# Super-Resolution Optical Sectioning Microscopy Visualizes Nanopores in the Plasma Membrane of Endothelial Cells *in situ*

**DOI:** 10.64898/2026.08.07.743430

**Authors:** Jasmin C. Schürstedt-Seher, Henning Ortkrass, Annika Kiel, Sylvia M. Steinecker, Wolfgang Hübner, Angela Kralemann-Köhler, Laureen P. Helweg, Marcel Müller, Jakob Wessendorf, Femke Testroet, Friedemann Kiefer, Jan Schulte am Esch, Thomas Huser

## Abstract

The ultrastructure of endothelial cells (ECs) “*in situ*” is of great interest due to their involvement in many physiological processes. In some organs, these cells form transcellular pores or fenestrae, allowing for the rapid exchange of molecules between blood and interstitium. Despite their importance, no optical images of these dynamic morphological structures have yet been acquired *in situ*. Major obstacles to their in-situ imaging are the lack of specifical labels for fenestrae and their size well below the optical diffraction limit. Here, we report how we have overcome these challenges and managed to visualize the EC ultrastructure *in situ* in 25 µm thick liver sections. To enable this, a lipophilic, fluorescent membrane dye was infused into the portal vein of murine livers to stain the sinusoidal ECs before the organ was harvested. Tissue sections were subsequently imaged using a novel, super-resolution optical-sectioning structured illumination microscope (OS-SIM), providing approx. 170 nm spatial resolution with significantly faster image acquisition compared to confocal microscopy.

## 1. Introduction

The liver, the largest organ in the human body is composed of a complex macroscopic and microscopic architecture that supports its indispensable function to maintain physiological homeostasis^1^. Imaging the vascular structure of the human liver is particularly challenging because of the need to cover length scales across seven orders of magnitude (from the nanometer scale to the centimeter scale^2^) to fully assess the ultrastructure of the entire organ down to the subcellular scale and probe its physiological function. Within the liver lobules, specialized fenestrated sinusoidal endothelial cells (LSECs) line capillaries with an inner diameter of less than 10 µm^3,4^ that form the functional interface between the blood and hepatocytes. Hepatic sinusoids are an extremely challenging target for high-resolution 3D imaging because of their small and highly curved spatial structure.

To support its physiological function, the liver requires the continuous and rapid exchange of metabolites with the blood. As a special morphological feature to warrant high metabolic throughput, the plasma membrane of LSECs contains 1000s of nanosized pores with a diameter of 50-200 nm, which function as molecular sieves^5,6^. LSECs retain circulating blood cells in the vessels, whereas molecules, such as small molecule metabolites, pharmaceutical compounds, macromolecules, as well as small particles, such as lipoproteins, viruses, and nanoscale drug formulations can freely pass through this barrier. Despite their importance, very little is known about the dynamic structure and function of these nanopores, or fenestrations, as they are well below the optical resolution limit and there are no specific cell surface markers for these structures.

Up until recently, the three-dimensional nanoscale morphology of liver sinusoids could only be imaged and analyzed *in-situ* by scanning electron microscopy (SEM)^7^. This requires time- and labor-intensive preparation of liver sections, the breaking of frozen tissue slices to reveal the sinusoids, and the resulting images typically only provide a small glimpse of the inside of a sinusoid^6,8,9^. Furthermore, specific labeling of surface markers in SEM is limited to labeling with gold particles, which is inefficient and only provides contrast for one type of biomolecule of interest (e.g. membrane proteins). More recently, the combination of focused ion beam (FIB) milling and SEM imaging was used to determine the liver ultrastructure at a spatial resolution down to approximately 8 nm^10^. Here, the FIB serves two purposes: it is used to remove (“ablate”) a small layer of tissue by scanning a focused beam of gallium ions across the surface. The surface is then imaged by SEM, while a significantly lower intensity gallium ion beam is still present to compensate the negative charge which the electrons deposit to the tissue. This alleviates the need for a conductive coating and enables the seamless imaging of newly “excavated” tissue structures. While FIB-SEM is an extremely powerful way of imaging tissue ultrastructure, it is a very time-consuming process with an expensive instrument that is only available at highly specialized laboratories. The subsequent data analysis is also very demanding, because no specific labeling of biomolecules can be used to help facilitate this process, and the identification of structures has to rely entirely on morphological features. Lastly, since material is removed during the scanning process, such an experiment can only be conducted once for every specimen. Because of these high demands and the high spatial resolution obtained with FIB-SEM, current imaging experiments are limited to approximately 15 vertical layers of hepatocytes^10^.

In order to visualize the nanoporous structure of LSECs with fluorescence microscopy *in situ*, several significant challenges have to be overcome. One problem of imaging thick liver slices with fluorescence microscopy is the scattering of light by the very dense tissue structure. The high level of intrinsic autofluorescence from tissue also results in strong out-of-focus contributions to any detected fluorescence signal. While this can be reduced by confocal microscopy, it severely affects all widefield imaging methods. Light scattering can be reduced by optical clearing of the samples and refractive index matching with aqueous or organic solvents^11^. Both organic solvents as well as detergents applied during clearing in aqueous solutions, however, aim to efficiently extract lipids from the sample to reduce optical heterogeneity and, unfortunately, result in the destruction of the ultrastructure of the cells’ plasma membranes. Therefore, optical clearing of liver slices is not suitable for the visualization of nanopores within the plasma membrane.

In order to overcome out-of-focus contributions, several different fluorescence microscopy techniques have recently been developed that manage to reduce the out-of-focus signal by providing optical sectioning of the sample^12^. One such option is light sheet fluorescence microscopy (LSFM), where the excitation and emission path are usually perpendicular to each other. Excitation light is focused into a thin optical sheet which only excites fluorescence in a single plane with 300 – 600 nm thickness and therefore avoids out-of-focus illumination and signals. LSFM is well suited for imaging thick tissue samples but typically requires optical clearing of the samples with all its limitations as mentioned above^13^.

Another approach to reduce the out-of-focus contributions is wide-field optical sectioning microscopy (OS). A coarse striped illumination pattern (with a pattern periodicity of several microns) is projected into the sample, such that fluorescence is only excited in the bright, illuminated stripes. By laterally shifting this pattern during three subsequent exposures, the parts of the image that are in the focus of the objective lens can be extracted and the out-of-focus contributions are effectively rejected^14,15^. This OS method has the advantage that it provides much faster imaging with optical sectioning contrast compared to the tedious scanning approach of a confocal laser-scanning optical microscope^16,17^. It is furthermore more photon-efficient than confocal microscopy.

Here, we demonstrate the combination of optical sectioning based on structured illumination microscopy (SIM)^18–20^ with super-resolution optical microscopy (OS-SIM). A sinusoidal interference pattern with a spacing of about 760 nm at an excitation wavelength of 647 nm is projected into the sample plane and shifted laterally three times. Subsequently, the pattern is rotated twice by 60°, and again three phase images are acquired, resulting in nine raw images per vertical slice. Computationally processing these raw images results in the effective suppression of out-of-focus signals, which are not modulated by the striped excitation pattern by employing a tailored notch filter in the signal processing chain^21^. In order to obtain a 3D image stack, the sample is then axially translated through the focal plane of the objective lens. The image acquisition scheme is similar to super-resolution structured illumination microscopy (SR-SIM) and by applying SR-SIM based image reconstruction algorithms, the spatial resolution of the resulting image data can be similarly improved. However, due to the rather coarse pattern, the OS-SIM approach prioritizes optical sectioning quality over the enhancement of lateral resolution. Thus, OS-SIM image reconstruction provides both, optical sectioning capability with approx. 600 nm axial sectioning (comparable to confocal microscopy) and a modest lateral resolution improvement resulting in approx. 170 nm spatial resolution, i.e. an approx. 1.35x resolution improvement. In combination with the in-situ perfusion of a lipophilic membrane dye, this technique allowed us to visualize nanoholes in the membrane of liver sinusoidal endothelial cells *in situ*.

## 2. Results

### 2.1 Efficient fluorescent staining of vascular endothelium and imaging by optical sectioning microscopy

To visualize the vascular network of the liver *in situ* with optical fluorescence microscopy, the cells of interest need to be specifically labeled. A particular complication of this process is that to date no specific markers have been found that would allow for the fluorescent labeling of fenestrations in LSEC. Instead, the entire plasma membrane of the endothelial cells needs to be labeled as efficiently and as homogeneously as possible, so that fenestrations can be identified as non-stained dark, nano-sized holes against the bright background of the plasma membrane. This requires the development of a specific staining protocol that will only fluorescently stain the membrane of endothelial cells as efficiently as possible, without creating an overwhelming background from the stained plasma membrane of all the other cells that are also present in the liver. This pretty much directly dismisses whole mount staining protocols, because the direct incubation of the tissue by application of a non-specific lipophilic membrane dye to an intact liver slice results in staining and fluorescent signal from all plasma and cytoplasmic membranes of all cell types present. To avoid this and to achieve specific staining of the endothelial cells, we perfused the dye directly into the portal vein of a recently sacrifized mouse to limit the staining process to cells in close contact with the blood i.e. the endothelial cells surrounding the blood vessels. This process and the choice of fluorescent stain, however, created another challenge: a lipophilic fluorophore embeds itself into the plasma membrane and is, thus, not fixed to a specific location. It will continue to diffuse within this environment. Naturally, this leads to leakage of the dye, which diffuses through the endothelium and eventually reaches the hepatocytes. The only way to overcome this challenge is to image the freshly perfused tissue as soon as possible post-perfusion.

The result of this approach is summarized in Figure 1. Here, maximum intensity projection micrographs of the same region of interest of a 20 µm thick liver slice perfused with BioTracker 655 red cytoplasmic membrane dye imaged by three different optical sectioning microscopy methods are shown. Figure 1a shows the result of imaging this vessel-stained liver slice with a commercial confocal laser-scanning microscope (CLSM). Figure 1b shows the same region of interest imaged by widefield optical sectioning microscopy, based on structured light that utilizes a reconstruction algorithm introduced by Neil and colleaques^14,22^. The pairwise subtraction of phase images *I*_*i*_ and subsequent summing of these squared differences effectively removes out-of-focus signal:

**Figure 1:**
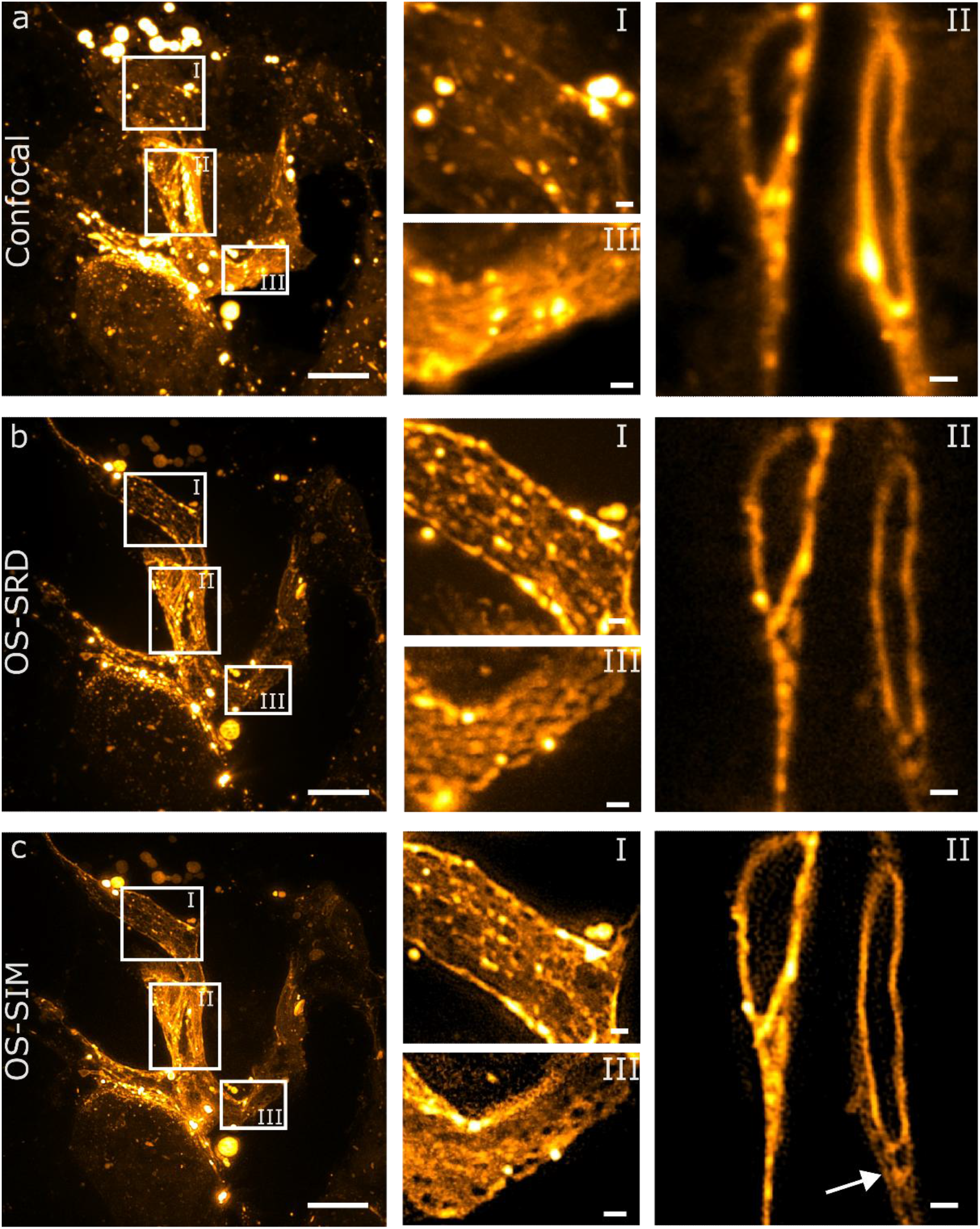
Comparison of imaging results of perfused vessel-stained mouse liver slices by confocal laser-scanning microscopy (CLSM), optical sectioning with the sum of reduced differences algorithm (OS-SRD), and OS-SIM imaging of the same region of interest. Maximum intensity projections of a 20 µm thick liver section imaged with (a) CLSM, (b) optical sectioning based on structured illumination but reconstructed with the sum of reduced differences algorithm (OS-SRD), and (c) OS-SIM. The cropped regions (I,III) show different areas of the vessel with small gaps/holes in the membrane structure. They are clearly resolvable in the OS-SIM image in contrast to the confocal microscopy image. In the hydrated tissue section, BioTracker 655 cytoplasmic membrane dye continues to diffuse to other cells and membranes. This is indicated by the cropped region (II) displaying a single z-slice of the stack where predominantly the nuclear membrane is visible but also the ER, vesicles and a low intensity plasma membrane (arrow). The pixel size of the confocal image was adjusted for better comparison and scaled to the same pixel size as that of the OS-SIM and OS-SRD images of 41.5 nm. The commercial confocal microscope was run in the “optimal” image scan mode provided by the manufacturer software, resulting in an original pixel size of 93.8 nm, which also limits the ability to compute FRC resolutions for this dataset, since at this pixel size, the confocal image is not Nyquist-sampled. Scale bar a-c: 10 µm; I-III: 1 µm.

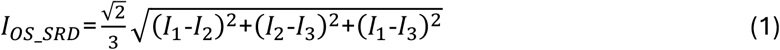

This reconstruction process suppresses the background and out-of-focus signal, but does not produce an enhanced resolution through frequency shifting, as obtained with the Gustafsson SR-SIM algorithm^19^. In the following we will refer to optical sectioning images reconstructed with the Neil algorithm as “sum of reduced differences optical sectioning” (OS-SRD). In contrast, the resolution-enhanced optical sectioning images will be referred to as OS-SIM. In order to permit both, OS-SRD and OS-SIM, we acquired nine raw images per z-slice (three angles, each with three phases). For reconstruction with the OS-SRD algorithm the three phase images for every angle were summed according to equation 1 and averaged over the three angles^23^. Figure 1c shows the OS-SIM reconstruction of the same region of interest by reconstructing the nine raw images using the SR-SIM reconstruction algorithm as implemented in the ImageJ-plugin fair-SIM^24^.

In practice, the CLSM image data (Fig. 1a) were acquired after the acquisition of the widefield optical sectioning image data was completed. This explains the presence of the bright dots of the dye that most prominently appear in the CLSM images which are due to the continued diffusion of the plasma membrane dye. Over time, this dye continues to diffuse through the tissue slice and accumulates in large dots, which are most likely lipid-rich vesicles. This also explains why the region of interest in Figure 1a I does not show the same intensity of the vessel staining as in the OS-SRD and OS-SIM images. Remarkably, both, the OS-SRD and OS-SIM images in the insets in Figures 1I and III reveal transcellular dark holes / pores with a diameter of 200-300 nm, which are not well resolved in the OS-SRD images. Figure 1II shows a single z-slice of the stack. From these images, it becomes apparent that the BioTracker 655 red cytoplasmic membrane dye also stains the nuclear membrane, the membrane of the endoplasmic reticulum (ER), and some vesicles. This provides evidence that this staining process is well suited to reveal cellular nanopores in endothelial cells *in situ*.

### 2.2 OS-SIM compared to confocal microscopy across a large field of view

To further explore how well the combination of vessel staining and OS-SIM imaging can resolve subwavelength structures in the liver vasculature, we then visualized a 25 µm thick liver slice across a larger field-of-view. Initially, a region of the tissue slice was chosen and mapped with our OS-SIM setup. Afterwards the same region was re-identified on the confocal microscope for correlative imaging. A 40x 1.1 NA water-immersion objective lens was used for confocal imaging to achieve the same field of view on both microscopes. This 40x objective lens allowed us to image and reconstruct an exceptionally large field of view by OS-SIM, but it sacrifices some of the resolution-improvement possible by OS-SIM due to the lower NA compared to that of an oil-immersion objective lens used for the OS-SIM imaging. The step size of the z-stack was set to 200 nm for both imaging systems. The correlative imaging results are shown in Figure 2. The maximum intensity projection and a single reconstructed z-slice are depicted in the first row of Figure 2 imaged by OS-SIM, while the second row shows corresponding regions imaged by CLSM.

**Figure 2:**
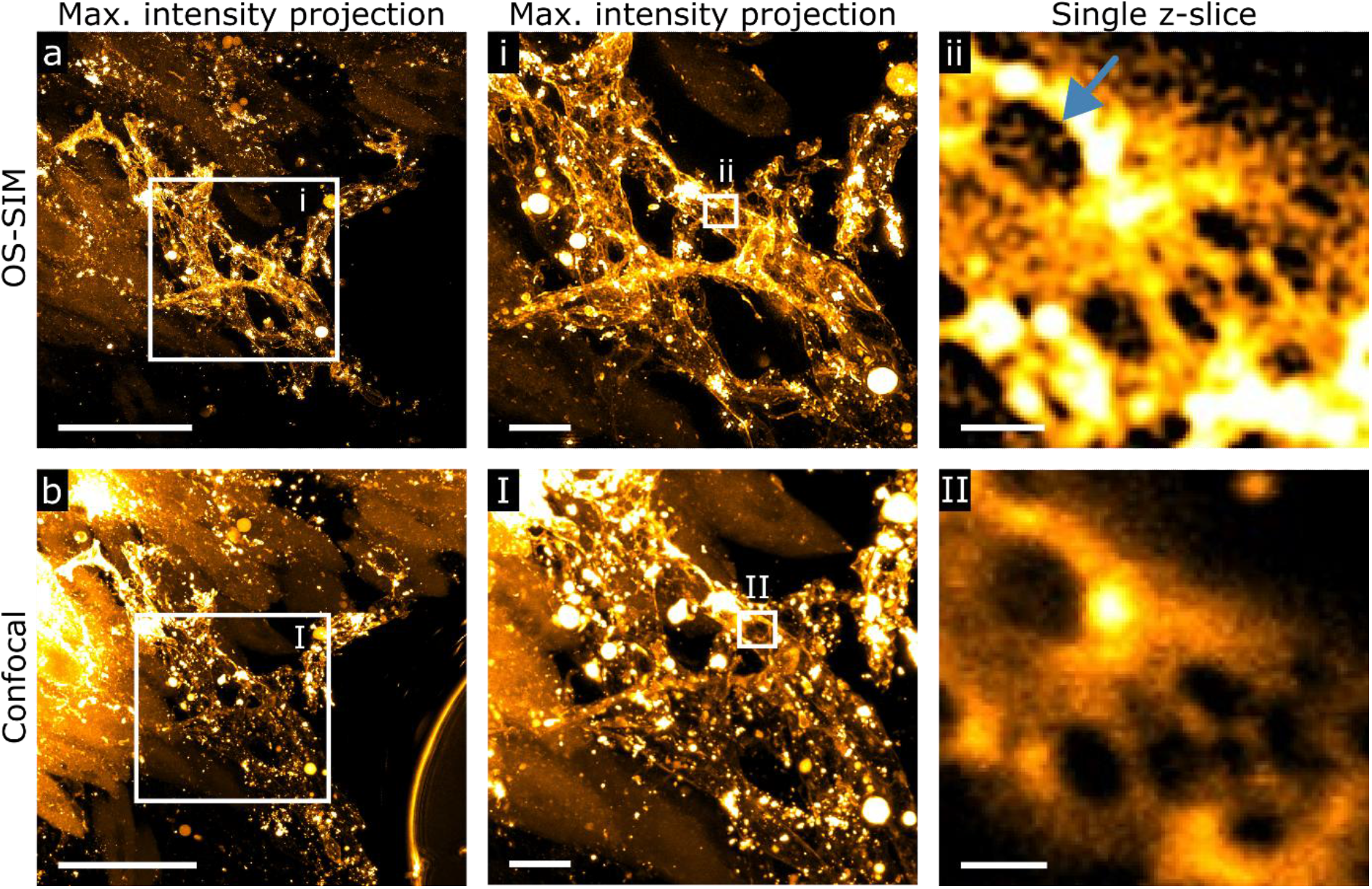
Comparison of confocal and OS-SIM fluorescence images of blood vessels in 25 µm thick liver slices perfused with BioTracker 555 Cytoplasmic Membrane Dye. Top row: OS-SIM reconstruction of the full 25 µm thick slice shown as (a) maximum intensity projection for a 162 µm x 162 µm FOV, alongside (i) a region of interest of the maximum intensity projection image and (ii) a region of interest from (i) shown as a single reconstructed z-slice. Bottom row: confocal laser-scanning fluorescence microscopy images of the same region. (b) Maximum intensity projection of the entire z-stack, (I) region of interest, and (II) single z-slice of the region of interest highlighted in (I). The overview images (a, b) present the vessel network but also reveal significant bleed-through of the perfused membrane dye to the hepatocytes, which is more prominent in the confocal image (b). The OS-SIM images reveal substructures inside larger holes in the vessel membrane reminiscent of a sieve plate (blue arrow) in a liver sinusoidal endothelial cell (ii). Scale bar a, b: 50 µm; i, I: 10 µm; I, II: 1 µm.

Figure 2a and b show a field of view of 162 x 162 µm, which allows for the visualization of an entire network of vessels. Bleed-through of the membrane dye to lower tissue structures (hepatocytes) is more prominent in the confocal image (Figure 2b). Here, the membrane of the surrounding hepatocytes in the upper left part of the image was also stained and appeared much brighter than the stained endothelial cell membrane in the middle. In the lower right corner of this image the mounting medium-air interface is visible, which does, however, not affect the image quality. This interface appeared over time in the mounting medium of the liver slice and is only visible in the confocal image because it was imaged after acquiring the OS-SIM data. The vessel network is even more clearly visible in the cropped region of the maximum intensity projection (Figure 2i, I). Note that the staining is not perfectly homogenous as there are some brighter fluorescent dots visible. These dots appear in different positions in the OS-SIM reconstruction and the confocal image, which is most likely due to the continued diffusion of the dye through the tissue. In the single z-slice (Figure 2ii, II) with the highest magnification the membrane of the endothelium surrounding the vessel is shown. Here, larger holes with a diameter of ∼1 µm can be seen in the plasma membrane. In the OS-SIM image the larger dark structures do not seem to be completely opened but appear to consist of several smaller holes with a diameter of 200-250 nm. These structures are reminiscent of sieve plates in LSEC^6^and were not resolved in the confocal image (Figure 2II).

### 2.3 Direct vessel painting stains specifically the endothelial cells in liver tissue

As seen in Figure 1, the cytoplasmic membrane dye BioTracker stains all plasma and internal cell organelle membranes. To further verify that perfusing the entire mouse liver with a membrane dye through the portal vein mostly stains the endothelial cells we conducted additional perfusion experiments with multiple fluorophores simultaneously. In addition to BioTracker 555, we perfused the liver with anti-PECAM-1 antibody direct-labeled with Alexa Fluor 647, an endothelial cell marker at the inner cell surface and more prominent at the cell junctions^25^. Furthermore, Hoechst 33342 was added to stain the nuclei of the tissue for reference. The result of this triple-staining experiment is shown in Figure 3. Figure 3a is a maximum-intensity OS-SIM image reconstruction of the BioTracker 555 channel. Figure 3b shows the same region of interest for the anti-PECAM1 channel and Figure 3c shows the Hoechst 33342 channel. Figure 3d depicts the overlay of the three different color channels as maximum intensity projection. Remarkably, the staining pattern of anti-PECAM1 and BioTracker 555 (Figs. 3a and 3b) is very similar, which indicates that BioTracker 555 initially stains the endothelial cells very well and quite selectively before diffusing into other areas of the liver tissue. The fluorescently labeled anti-PECAM1 shows a higher signal at the circumference of the endothelial cells as expected (see Fig. 3b). However, due to the direct perfusion of the antibody without extra washing and blocking steps, other parts of the endothelial cells are also stained. The same cells are also very homogeneously stained with the BioTracker 555 dye (Fig. 3a), which confirms that the cytoplasmic membrane dye is a good choice in order to identify and image fenestrations in these endothelial cells. The nuclei of both, the hepatocytes and the endothelial cells are labeled by Hoechst 33342 as shown in Figure 3c. The Hoechst channel also reveals that the nuclei of the hepatocytes have a typical round shape, whereas the nucleus of the endothelial cells is more elongated, which provides another means of identifying and separating these cells in-situ (see Figure S3).

**Figure 3:**
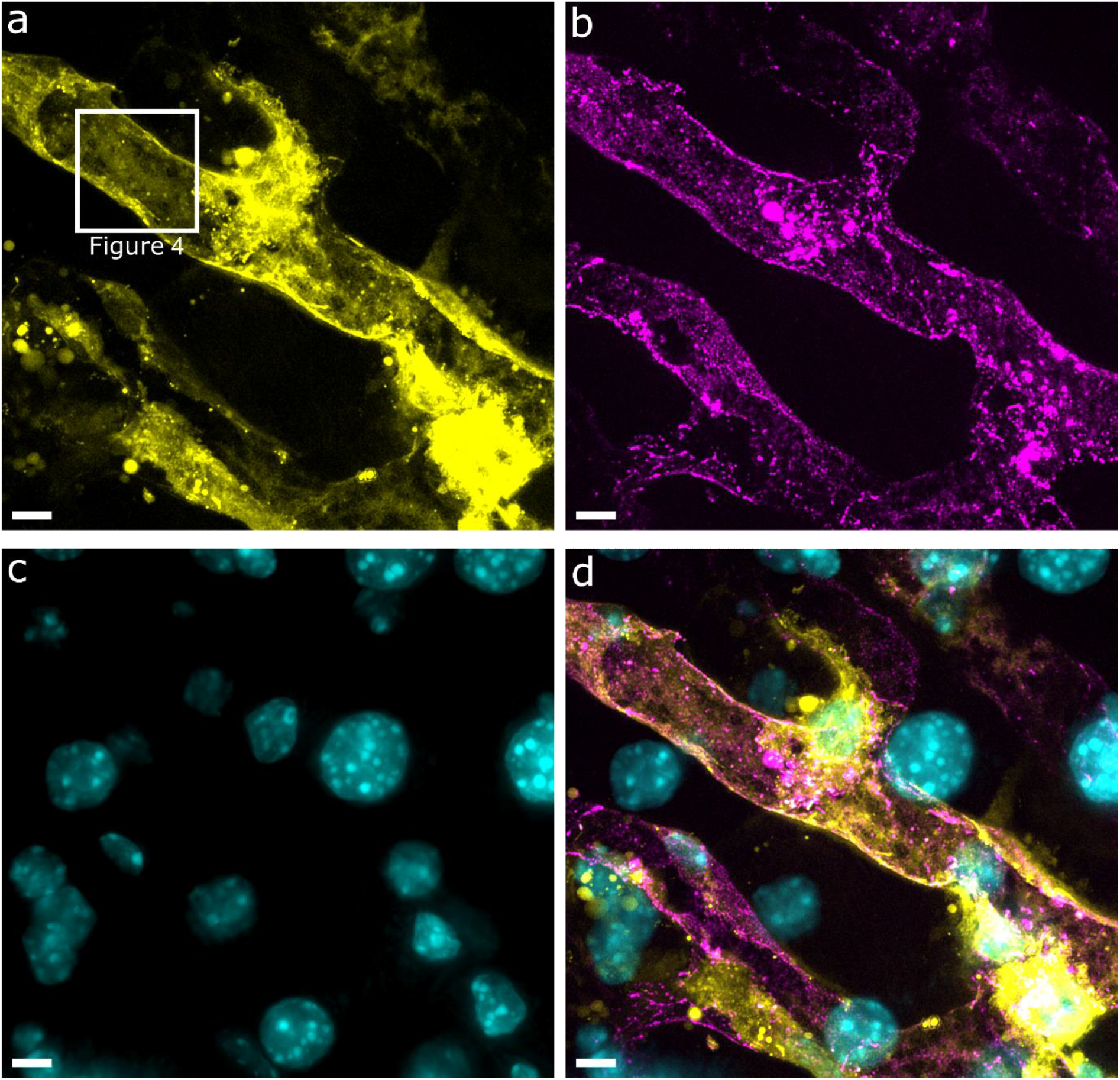
OS-SIM reconstruction of a perfused liver tissue slice shows specific staining of endothelial cells lining the arteries. The mouse liver slice was perfused with (a) cytoplasmic membrane dye BioTracker 555 (shown in yellow), (b) anti-PECAM-1 antibody directly labeled with Alexa Fluor 647 (magenta), and (c) Hoechst 33342 (cyan). (d) Overlay of all channels. The maximum intensity projection of a 14 µm OS-SIM reconstructed z-stack is shown in (a) and (b) and the widefield image projection for the Hoechst channel is visible in (c). Based on the colocalization of the stains it is apparent that the endothelial cells surrounding the blood vessel are stained with both, the membrane dye (a), and the specific endothelial marker PECAM-1 (b). The nuclei of the endothelial cells as well as the surrounding hepatocytes are stained with Hoechst (c), but contrast for the nuclei of the endothelial cells is reduced due to their shape and smaller volume . Scale bar a – d: 5 µm.

Close inspection of Figure 3a reveals that there are multiple areas, where groups of dark holes can be identified within the fluorescent plasma membrane. Magnified views of the areas highlighted by white outlines in Figure 3a are shown in Figure 4. The size and grouping of these apparent nanoholes is, yet again, indicative of sieve plates in LSEC.

**Figure 4:**
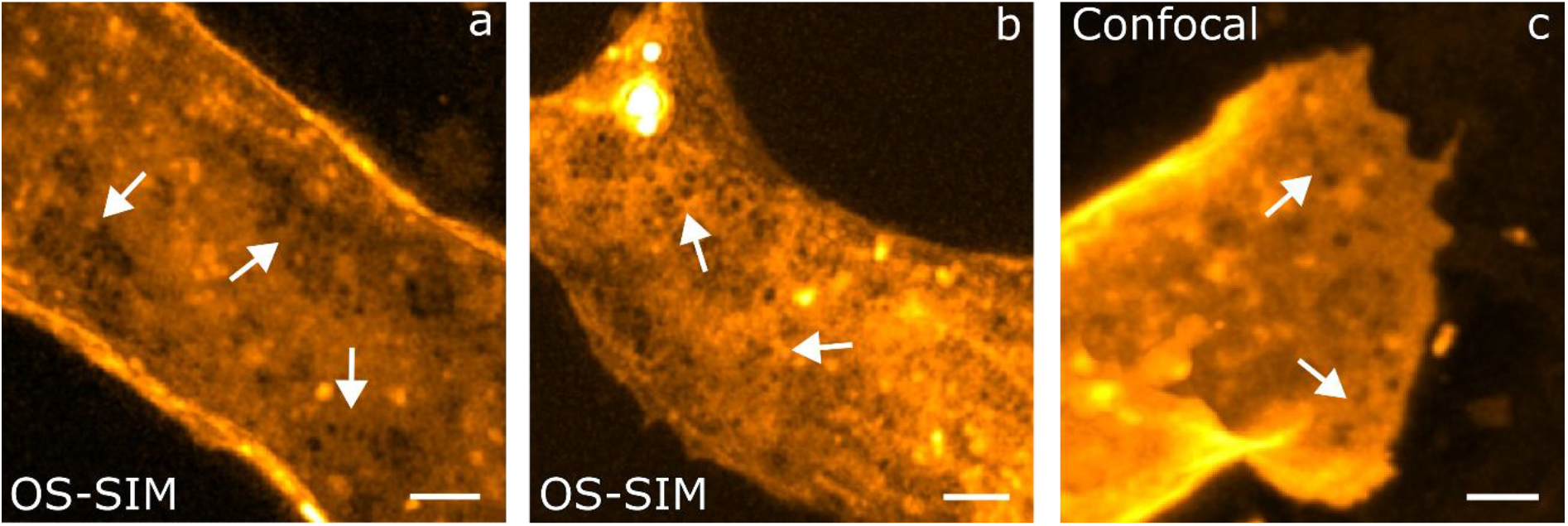
OS-SIM is able to resolve entire sieve plates in-situ in liver sinusoidal endothelial cells. For better visualization a 1-pixel Gaussian blur is applied to both, the OS-SIM and confocal microscopy images. (a, b) Magnified views of regions of interest highlighted by a white outline in Fig. 3a. The images are maximum intensity projections of one half of a stained vessel (perfused with BioTracker 555) of a OS-SIM image reconstruction, which better visualizes the sieve plates within the endothelial membrane. (c) A comparable position of a perfused liver slice imaged by CLSM. The arrows point to regions indicative of sieve plates. Scale bar a-c: 2 µm.

### 2.4 Quantitative analysis of the resolution improvement achieved by OS-SIM

In order to more quantitatively determine the spatial resolution that can be achieved by OS-SIM, we applied unbiased resolution measurements to both, OS-SIM reconstructed images of perfused blood vessels and widefield images reconstructed from the same raw images. Figure 5a shows the maximum intensity projection of OS-SIM and widefield fluorescence images of a 25 µm thick mouse liver slice perfused with BioTracker 655 red cytoplasmic membrane dye. Here, a 40x objective lens with an excitation laser wavelength of 647 nm was used for imaging. The filtered widefield image is the average sum of all nine raw images per z-slice, where an additional Wiener filter was applied. The upper left part of the vessel is outside the imaged depth range and therefore not visible in the projection. The holes with a diameter > 300 nm in the membrane of the endothelium are faithfully reconstructed without any artifacts (Figure 5i) and appear blurrier in the filtered widefield image (Figure 5I). The smaller holes with a diameter of 200-300 nm in the region of interest in Figure 5ii are not visible in the filtered widefield image (Figure 5II). The analysis of the achieved resolution was performed by Fourier ring correlation (FRC)^26^ (Figure 5c), which requires two independent images of the same region for the correlation analysis. Because the raw data were only acquired once, we used an algorithm to coin-flip the data set into two independent sets^27^. The photon counts for each pixel of the raw data stack was randomly distributed into two separate stacks. These stacks were individually reconstructed using the same parameters in fairSIM, resulting in two independent reconstructed stacks and the corresponding pseudo widefield (sum of all raw images per slice) and filtered widefield images with additional Wiener filtering for visualization. The FRC analysis was done on one slice of the z-stack close to the coverslip surface in order to calculate the best resolution achieved, as shown in Figure 5c.

**Figure 5:**
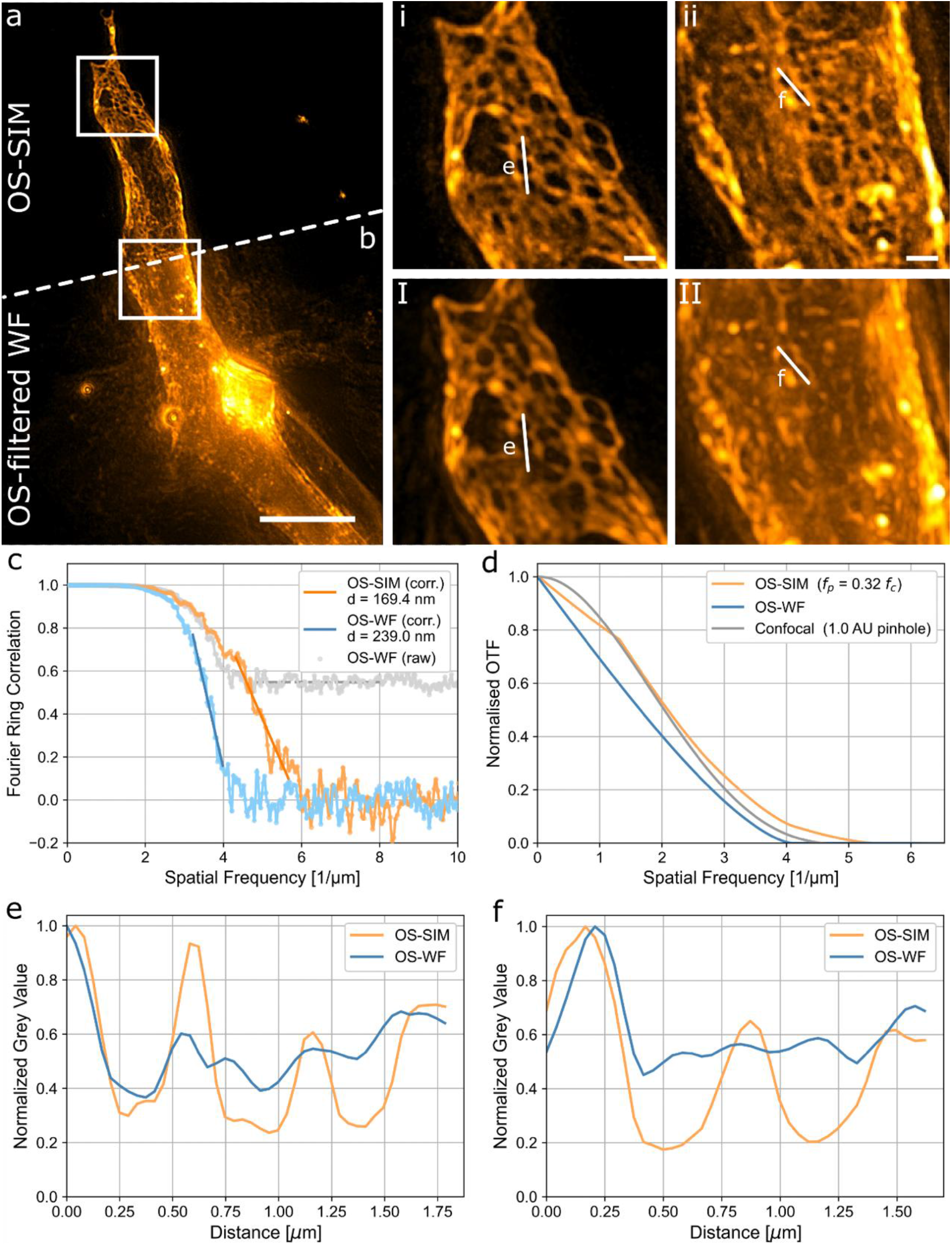
Comparison of a reconstructed OS-SIM image and the corresponding filtered widefield fluorescence image of a 25 µm thick liver slice stained with BioTracker 655 Cytoplasmic Membrane Dye. (a) Upper half: maximum intensity projection of the reconstructed 25 µm OS-SIM z-stack. (b) Lower half: filtered widefield image. The regions of interest of the OS-SIM images are presented in(i), (ii) and the corresponding filtered widefield images in (I), (II). The OS-SIM images (i) and (ii) clearly reveal distinct nanopores in the fluorescently labeled plasma membrane, whereas the same structures appear very blurry in the filtered widefield images (I), (II). (c) Fourier ring correlation (FRC) plots indicate an enhanced spatial resolution of the OS-SIM (orange) image of 170 nm compared to the OS-WF (blue) image with 240 nm. (d) Theoretical support of the effective optical transfer function (OTF) for the different imaging modalities. The pattern spacing of the OS-SIM of the calculated OTF was set to 760 nm, corresponding to the real pattern spacing for imaging. The OTF of the OS-SIM (orange) is above the OS-WF (blue) and confocal (gray) indicating the higher frequency support in the low to high frequency range. (e, f) line plots corresponding to the white lines in (i, ii) and (I, II). Scalebar a, b: 10 µm; i, ii, I, II: 1µm

The FRC curve of the widefield and reconstructed coin-flipped images showed an unphysical, too high correlation in the high frequency range (OS-WF raw, gray plot in Figure 5c). This is a well-known effect caused by correlations introduced by fixed noise pattern of the camera^28,29^. We used an established process to correct the FRC curve, where the mean of the baseline at the high correlation plateau was fitted and subtracted. Afterwards the data were rescaled to a maximum correlation of 1 and the linear slope of the curves was fitted (Figure 5c orange: OS-SIM corr. and blue: OS-WF corr.). To estimate the spatial resolution, the zero of the fit function was calculated. The result shows that OS-SIM achieves a spatial resolution of 170 nm compared to the resolution of the pseudo widefield image of ∼240 nm. Both, the resolution limit and the FRC curve shape fits well with the effective optical transfer functions (OTF) for both widefield and OS-SIM, that are presented in Figure 5d. The frequency support for the OS-SIM with a pattern spacing of 760 nm, that was used for acquiring the data, is higher for the low to high frequency range compared to the classical widefield and even for the confocal. The lineplots in Figure 5e and 5f visualize this behavior of the higher frequency support and the improved resolution of the OS-SIM imaging. OS-SIM achieves a higher contrast in the resolved structures and even the small holes in the membrane can be differentiated in the OS-SIM image whereas the filtered widefield image only allows a visualization of the larger holes.

## 3. Discussion

Fenestrations in LSECs with a diameter of less than 200 nm are typically grouped together in sieve plates, where the individual fenestrae are separated by actin filaments and sieve plates are surrounded by tubulin^30,31^. In cultured LSECs it has been shown that the thickness of the entire cell (from the basal to the apical plasma membrane) in the region of sieve plates is usually less than 200 nm^32^, resulting in an extremely thin and highly porous barrier between blood and the organ. In cultured cells the staining of the plasma and cytoplasmic membranes lets the sieve plate region appear less bright in super-resolution fluorescence microscopy images of LSEC because of the reduced thickness of the cell^33–35^. We speculate that the larger holes in the plasma membrane of the cell layer in the lumen of arteries as shown in Figure 2 could represent sieve plates. Clearly, the spatial resolution of the confocal microscope is not sufficient to resolve individual fenestrations, but it shows larger dark areas that appear to be either big holes in the plasma membrane or unstained areas in the membrane of the endothelial cell layer. These dark areas are reminiscent of sieve plates with significantly lower fluorescence signal, where the fenestrations inside could not be resolved. The enhanced spatial resolution provided by OS-SIM allows us to resolve some smaller structures with a diameter of approximately 200 nm within the darker regions of the membrane (Figure 2ii). This observation supports our hypothesis that OS-SIM can potentially resolve individual fenestrations within the sieve plates of liver sinusoidal endothelial cells *in situ*. The size of the other resolvable small holes (Figure 4a&b, Figure 5a II) is in the same size range of about 250 nm and they appear grouped together, which matches with the reported size and arrangement of fenestrations^31^. These intracellular holes are present in all the reconstructed images acquired by OS-SIM independent of the plasma membrane stain (BioTracker 655 in Figure 1 and 5, BioTracker 555 in Figure 2, BioTracker NIR750 in Figure S1 and S2) used in the image acquisition, which indicates that these are not caused by labeling artefacts.

The big holes that appear in the endothelial cell plasma membrane in Figure 2ii could either be larger gaps between LSECs, transendothelial channels (TEC)^36^ or iGap formation (holes larger than 300 nm) due to inflammation^37^. Despite this uncertainty, the OS-SIM reconstruction of these bigger holes also reveals a substructure indicative of arrangements of nanoholes in the plasma membrane, which are even better resolved and more defined when stained with Bio-Tracker 555 (see Fig. 4a,b).

Lipophilic carbocyanine membrane dyes such as the BioTracker cytoplasmic membrane dyes used in this study do not only specifically stain the plasma membrane of the cells, but also other cell membranes. The membranes of all internal cell structures are also mostly labelled.

The nuclear membrane and the mesh like structure of the ER and Golgi apparatus combined with mitochondria are visible in Figure 1. Interestingly, the plasma membrane here appears with significantly lower brightness.

In the NIR wavelength range (785 nm) the optical sectioning of the liver tissue works as well. The difference of the NIR OS-SIM reconstruction and the filtered widefield image shows the rejection of the out-of-focus signal in Figure S2. Here, the spatial resolution that we obtain is reduced by the longer excitation wavelength and therefore it is harder so resolve individual nanopores in the membrane of the endothelial cells. Nonetheless, it is possible to image areas containing small holes that could, again, indicate the presence of fenestrations (Figure S1,S2).

The imaging speed of our OS-SIM setup is much faster than that of commercial CLSMs since it is not a point scanning but a widefield approach. Although 9 raw images are acquired in order to reconstruct a single z-slice, the acquisition time for these raw images is (dependent on the exposure time, usually 50 ms) in the range of 1-2 s per z-slice. Typically, the acquisition time for a z-stack imaged with a confocal microscope takes several minutes and not uncommon up to more than 10 minutes.

The limitation of this OS-SIM technique with enhanced resolution is the reconstruction of the images. It is based on the standard Gustafsson algorithm^19^ and therefore reconstruction artefacts can occur^38^. The classical optical sectioning algorithm first introduced by Neil based on the differences of the phase images is not prone to these image reconstruction artefacts but on the other hand is limited in the achieved spatial resolution. Due to the thick sample and differences in refractive indices, spherical aberration becomes visible especially in deeper lying areas that are further away from the objective lens, and these cannot be corrected. In Figure 2 the cropped region of the OS-SIM reconstruction shows some reconstruction artefacts due to modulated out-of-focus signal and spherical aberrations where the approximated OTF is not fitting resulting in spotty signal. This becomes visible at the edges of the vessel and deeper inside the tissue.

Another limitation is the tube-like structure of the vessels and the non-isotropic resolution of the OS-SIM reconstruction. We gain a lateral resolution enhancement but, in the z-direction the resolution is limited to the widefield resolution.

## 4. Methods

### 4.1 Optical Setup

Optical sectioning structured illumination microscopy (OS-SIM) data were acquired on two bespoke setups, one working in the visible range (using lasers at 405 nm, 561 nm and 647 nm excitation wavelength) and another working in the NIR wavelength range (with 785 nm laser excitation)^28^. The working principle of both setups is similar: a modified Michelson-type interferometer is used to generate the sinusoidal interference by two mutually coherent laser beams^39,40^. The position of the excitation laser in the back focal plane (BFP) can be set using a galvanometric mirror pair and the laser beam is split into two arms using a non-polarizing 50:50 beamsplitter cube. One arm of the interferometer is delayed by a glass window to achieve a phase difference.

By tilting the glass window at different angles, the optical path length of this interferometer arm is modified resulting in different phase steps. The other interferometer arm contains a retroreflector that reflects the beam back while mirroring it with respect to the optical axis. The distance of the beams to the optical axis and thus the excitation pattern spacing can be changed by the galvanometric mirror pair. The two beams are projected into the sample plane via a 4-f telescope where they interfere and generate the sinusoidal illumination pattern. Switching objective lenses between e.g. an UPlanXApo 40x 1.4 NA oil or a 60x Plan Apo 1.45 NA TIRF oil Olympus objective lens, allows us to vary the field of view. In the VIS setup two 500 mm achromatic lenses (ACT508-500-A, Thorlabs) arranged in the Ploessl configuration result in an effective focal length of 250 mm and act as tubelens in front of a pco.edge 10 bi sCMOS camera. For the NIR setup we used a 180 mm (U-SWATLU, Olympus) tube lens in front of a NIR-enhanced sCMOS camera (pco.edge 26 NIR). This results in an effective pixel size of 83 nm (40x, VIS) / 41.6 nm (60x, NIR) / 61.5 nm (40x, NIR). A schematic setup of the two interferometric OS-SIM systems is shown in Figure 6a.

**Figure 6:**
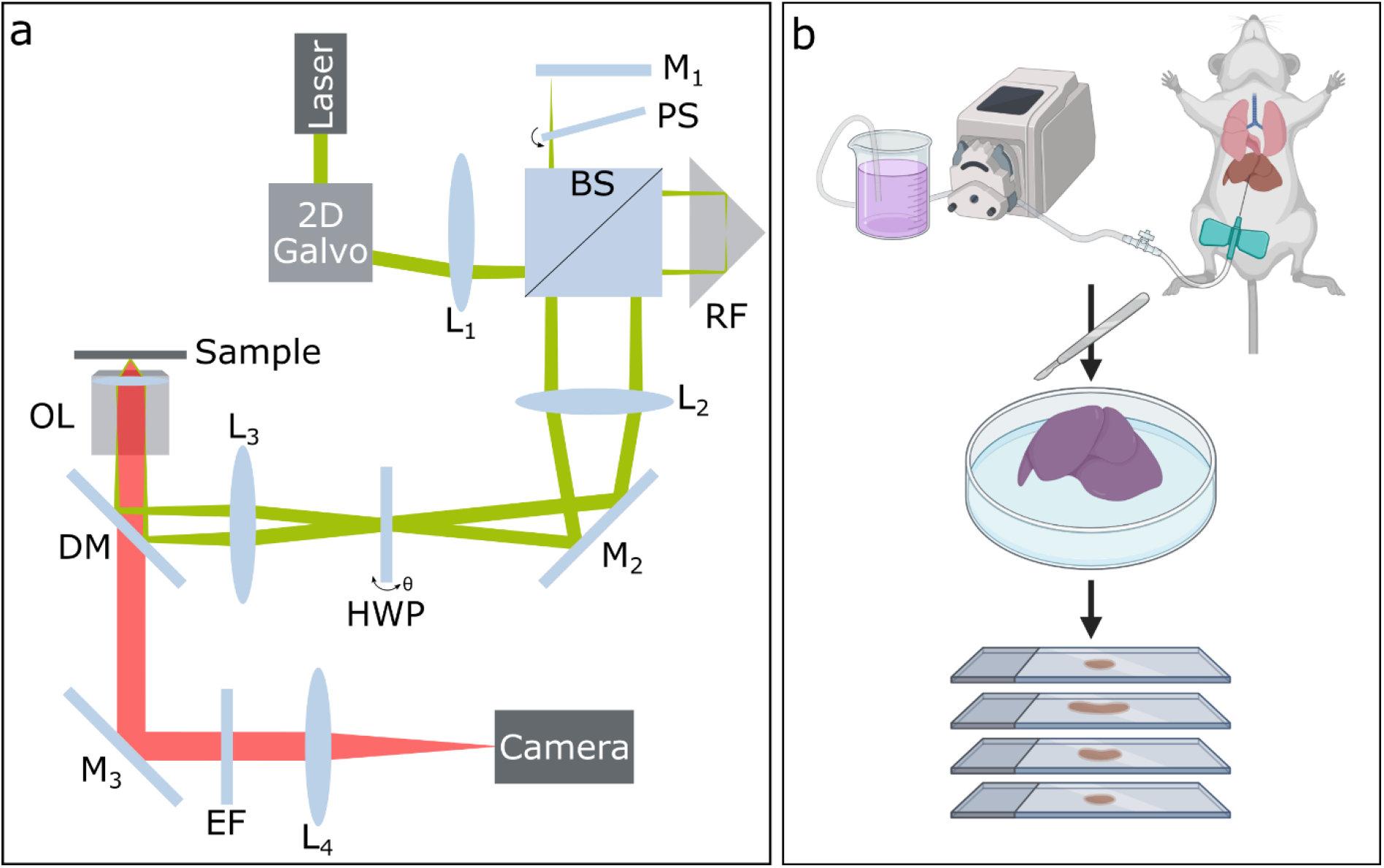
Schematic overview of the optical setup and sample preparation. The OS-SIM setup (a) is based on a modified Michelson-interferometer. The excitation laser beam (green) is reflected by a galvanometric mirror pair (2D galvo). The two beams are generated using a 50:50 beamsplitter cube (BS) where in one arm a phaseshifter (PS) is added to modulate the phase and the other arm consists of a retroreflector (RF) to flip the beam along the optical axis. The s-polarization of the beams is set using a rotating half wave plate (HWP). The beams are focused into the back-focal plane of the objective lens (OL) and the interference at the sample plane creats a striped illumination pattern. The fluorescence emission (red) is filtered through an emission filter (EF) and imaged onto the camera. L: lens, M: mirror, DM: dichroic mirror. For sample preparation (b) the liver of a mouse is perfused in-situ with the lipophilic dye, then removed and fixed overnight. After embedding in agarose gel it is sliced into 15-25 µm thick sections that are mounted on a microscope slide. b: Created with BioRender.com.

In order to provide the optical sectioning capability needed to image up to 25 µm thick tissue samples, the pattern spacing of the sinusoidal illumination needs to be coarser than for regular super-resolution structured illumination microscopy (typically around 200-250 nm). Scattering and absorption of the tissue distorts the projection of the pattern into the sample plane and result, together with high background signal, in a loss of visibility of the pattern, which is lessened by using a coarser pattern. For the VIS we used a pattern spacing of 760 nm and for the NIR samples the pattern spacing was varied from 610 nm with the 60x objective lens to 910-950 nm for the 40x objective lens.

We acquired z-stacks with a z-spacing of 200 nm across entire murine liver samples cut to 15-25 µm thickness with a vibratome. For every optical slice, nine raw images (three angles, each with three phases) are acquired for the VIS, whereas for the NIR we acquired 15 raw images (three angles, each with five phases) per z-slice. The images are reconstructed using the FIJI plugin fairSIM^24^ where the phase steps are fitted individually. It also generates a pseudo wide-field image as the sum of the raw images and a filtered widefield image with additional Wiener filter that can be used for comparison of image quality and resolution.

### 4.2 Sample Preparation

#### Animals

RjHan:NMRI mice, an in-house breeding line from the Animal Facility, faculty of Biology at Bielefeld University and *Vegfr3*-tdTomato transgenic mice^41^ from the Animal Facility at University of Münster were used. Mice were kept under standard conditions with water and chow (SSniff, regular chow diet) ad libitum. The mice used for the experiments were between 9 and 22 months old. Animals were euthanized by cervical dislocation. All animal experiments were approved by the respective local authorities and were in accordance with institutional guidelines for the welfare of animals.

#### Perfusion of the membrane dye

RjHan:NMRI or *Vegfr3*-tdTomato transgenic mice were sacrificed, the abdomen was opened and a cannula was carefully inserted through a small incision into the portal vein avoiding rupture of the vessel. Subsequently, the liver was perfused with buffer^42^ at a flow rate of 7-10 ml/min until blood no further exited the organ. Then the perfusion was continued with a dilution of lipophilic, fluorescent membrane dye in phosphate-buffered saline (PBS) until the liver exhibited a noticeable color change due to absorption of the dye (“Vesselpaint”^43^). The dilution was dependent on the membrane dyes applied, we used 1:1000 dilution for BioTracker 555 orange (SCT107, Sigma) and BioTracker 655 red (SCT108, Sigma) and 1:200 dilution for BioTracker NIR790 (SCT115, Sigma). For the multicolor staining we used a slightly different concentration of 1:200 dilution for BioTracker 555, 1:1000 dilution of direct labeled rat monoclonal anti-mouse PECAM-1 antibody (clones 5D2.6 and 1G5.1)^44^ with Alexa Fluor 647 and 1:10000 dilution for Hoechst 33258 (H1398, ThermoFisher). At the end of the procedure, the cannula was removed from the portal vein and the liver was excised and fixed using 4% formaldehyde (FA) in PBS over night at 4°C.

The fixed liver was embedded in 5% low melting point agarose and cut in 15-25 µm thick slices using a Leica VT1000S vibratome. The slices were stored in PBS until they were used for imaging. A schematic overview of the sample preparation is shown in Figure 6b.

#### Slice preparation for imaging

The liver slices only perfused with the BioTracker dye were embedded in ProLong Glass Antifade Mountant with NucBlue (P36985, ThermoFisher) on a glass slide, covered with a #1.5 coverslip and dried overnight. The liver slice with the multicolor staining was embedded in Vectashield (Vector Laboratories), covered with the coverslip and imaged directly. The mounted slices were imaged with the OS-SIM setup. For comparison some samples were also imaged with a commercial STELLARIS confocal microscope (Leica). We used a 40x 1.1 NA water objective lens to achieve the same field of view for both imaging techniques (Figure 2). To allow for a better comparison of the spatial resolutions, we used a 100x 1.4 NA oil objective lens (Figure 1). This objective lens was chosen because it has the same numerical aperture as the 40x objective lens used for VIS OS-SIM imaging. The step size for acquiring the z-stack was the same as for OS-SIM imaging and set to 200 nm.

Unfortunately, the native fluorescence signal from the transgenic mice was too weak in the liver sections so that no data could be recorded in the tdTomato channel.

## 5. Conclusions

We have overcome multiple challenges to demonstrate the visualization of nanopores in the membrane of liver sinusoidal endothelial cells *in situ* in 25 µm liver sections. These structures have so far evaded their visualization with fluorescence microscopy *in situ*. One challenge is the “negative” staining of these nanopores which have a diameter below the diffraction limit of light (< 250 nm). If the membrane of LSECs is stained, fenestrations appear as dark areas inside the membrane. To resolve these structures, super resolution microscopy is necessary. Another challenge with imaging thicker tissue sections is the scattering and absorption of the tissue that alters the image quality and resolution. With our OS-SIM method, we gain a moderate lateral resolution enhancement (170 nm) while also reducing the unwanted out-of-focus signal due to a lateral structuring of the illumination light with a periodicity near the wavelength of the excitation light. The endothelial layer inside the arteries of the liver section are stained by perfusing the fresh mouse liver with a warm phosphate-buffered saline solution containing the lipophilic membrane dye (BioTracker 555 orange, BioTracker 655 red or BioTracker NIR790 cytoplasmic membrane dyes). This treatment results in efficient fluorescent staining of the vascular endothelium. The moderately enhanced spatial resolution obtained with our OS-SIM setup readily allows us to identify structures reminiscent of sieve plates containing several cellular nanopores. This method opens up new possibilities for the investigation of morphological alterations of the tiniest hepatic arteries as a result of different diseases and disease progression.

## Acknowledgements

This project received funding from the Deutsche Forschungsgemeinschaft (DFG), grant number 540217954 and CRC1450–431460824. Part of the project was supported by the European Union’s European Innovation Council (EIC) PATHFINDER Open Programme under grant agreement No.101046928. This project was also supported in parts by the German Federal Ministry for Research, Technology and Aeronautics (BMFTR), through project BetterView (FKZ 13N15827). This project has also received funding from the European Regional Development Fund (EFRE/JTF-Programm NRW 2021-2027), funding line “Start-up Transfer.NRW” under the project “ProSIM”, as well as from the EXIST-Research Transfer program (Phase I) of the German Federal Ministry for Economic Affairs and Energy (grant number 03EFNW0436).

The authors would like to thank Karolina Szafranska for providing the idea of the perfusion of the liver *in situ*.

## Author contributions

J.C.S.S. performed the measurements of the samples on the OS-SIM and confocal microscope, reconstructed the data, analyzed the data and prepared the manuscript. H.O. built the interferometer-based SIM microscope that was used for the VIS measurements. A.K. initialized the liver perfusion of the mice and performed some of the sample preparations together with A.K.K. W.H. acquired the confocal imaging data of the slices. S.M.S. built the NIR SIM setup and conducted the NIR OS-SIM measurements. M.M. calculated the effective optical transfer functions. J.W. wrote the code for optical sectioning based on the sum of reduced differences algorithm. F.T. and F.K. provided the transgenic mice and discussed the protocols. J.S.a.E., F.K., and T.H. funded and supervised the work. All authors reviewed and agreed to the final manuscript.

## Data availability statement

The raw data is available on zenodo: 10.5281/zenodo.18798098.

