## Supplemental Material for "Super-Resolution Optical Sectioning Microscopy Visualizes Nanopores in the Plasma Membrane of Endothelial Cells *in situ*"

### 1. OS-SIM imaging in the NIR wavelength range

In the NIR wavelength range the penetration depth in tissue is increased compared to the visible spectral range (400-700 nm) because of the reduced scattering and absorption of the tissue[1,2]. We recently developed a NIR-SIM setup[3] were the pattern spacing of the sinusoidal illumination can also be adjusted to allow for imaging 25 µm thick liver tissue sections and visualize the membrane structure of endothelial cells. Here, the mouse liver was perfused with BioTracker NIR790 cytoplasmic membrane dye and fluorescence was excited with a 785 nm diode laser. 15 raw images were captured per z-slice (three angles, each with five phases) and reconstructed using fairSIM. The higher number of slices provides enhanced signal-to-noise ratio in cases where there is random phase drift, as the raw phase for every image is computed and compensated in the reconstruction[4]. Figure S1 shows the NIR OS-SIM reconstructions of two separate fluorescently stained samples. The maximum intensity projection image shown in Figure S1a was acquired with a 40x objective lens. The cropped regions show the plasma membrane with small pores (Figure S1I) and the nuclear and ER membrane (Figure S1II) of the first layer of cells lining the arterial lumen. The staining of the vessel in the maximum intensity projection of Figure S1b (imaged with the 60x objective lens) appears spottier. Some transcellular holes are visible in the single z-slice of Figure S1III and pores with a diameter of 270-410 nm are visible in the cropped region in Figure S1IV.

Figure S2 details the improvement in image quality of the NIR OS-SIM reconstruction in comparison to the filtered widefield image. Here, the maximum intensity projection of a 10 µm thick z-stack of a liver slice perfused with BioTracker NIR790 cytoplasmic membrane dye is shown. The first row represents the NIR OS-SIM reconstructed images, and the second row shows the corresponding filtered widefield images. The background signal is almost entirely suppressed in the NIR OS-SIM (Figure S2a) when compared to the filtered widefield image in Figure S2b. The difference is even more prominent in the cropped region where small intracellular holes with a diameter of 210 nm are visible in the membrane of the endothelial cells whereas the scattering of the tissue in the filtered widefield completely blocks the visualization of the membrane structure (Figure S2i and I) and results in noise artefacts. The nuclear membrane in Figure S2ii is better resolved as well when compared to the filtered widefield image.

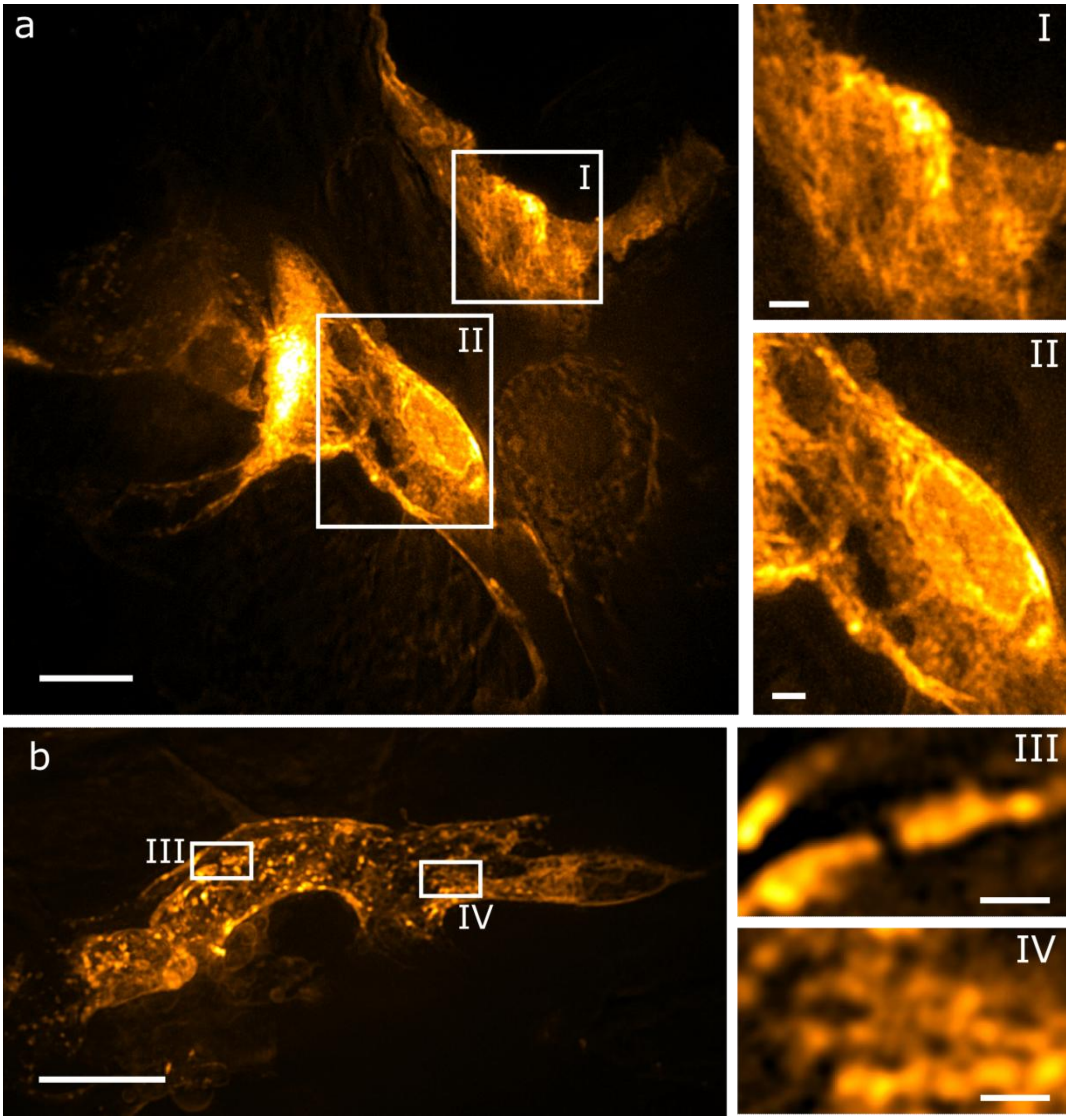


***Figure S1: NIR OS-SIM reconstruction of 25 µm liver slices perfused in-situ with BioTracker NIR790 cytoplasmic membrane dye.*** *The maximum intensity projection of a 10 µm z-stack imaged with a 40x objective lens in the NIR is shown in a. The corresponding regions of interest show the plasma membrane staining revealing small holes (I) and the staining of the nuclear membrane and ER structures (II). In b the maximum intensity projection of a 10 µm z-stack of a liver slice imaged with a 60x objective lens in the NIR is presented. The insets show single z-slices where some gaps in the membrane are visible (III) and also small holes (IV). Scale bar a, b: 10 µm; I,II: 2 µm; III,IV: 1 µm.*

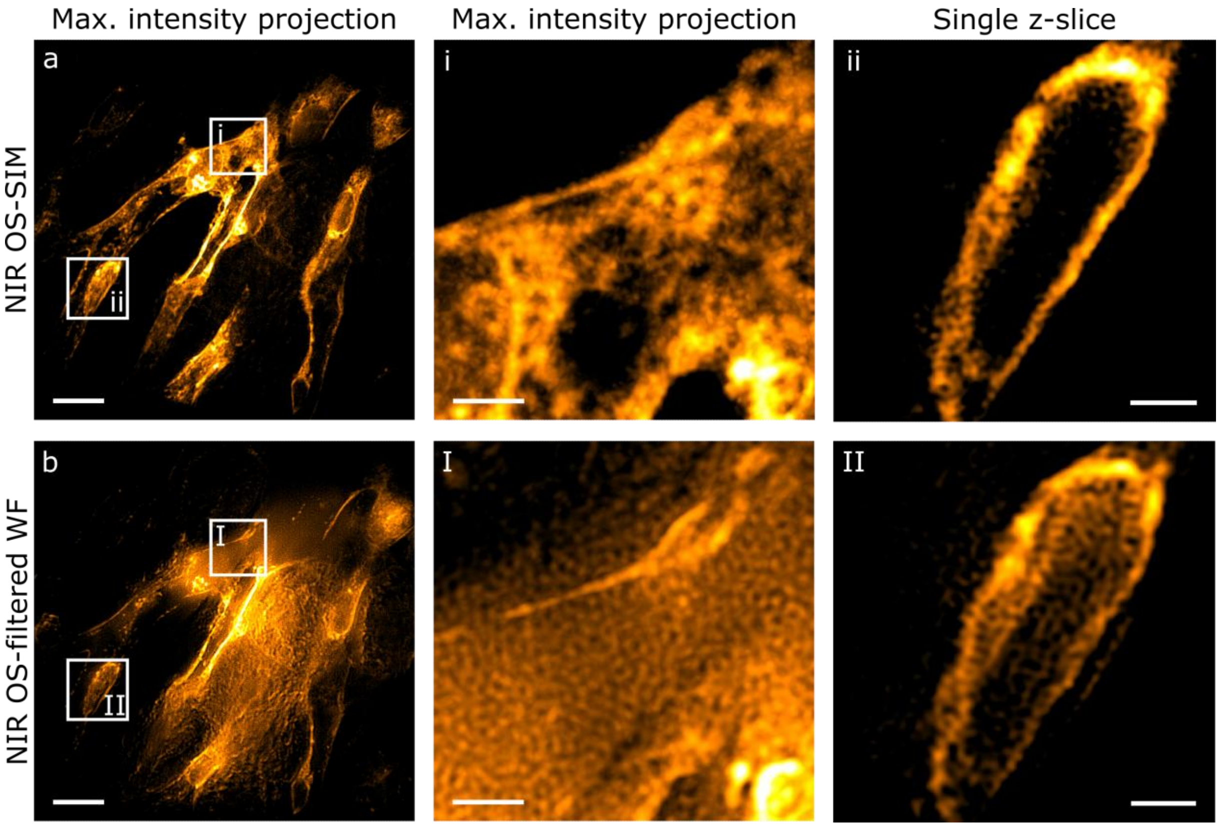


***Figure S2: Comparison of a reconstructed NIR OS-SIM image and the corresponding filtered widefield image generated with fairSIM.*** *The first row of the figure shows the NIR OS-SIM reconstructions and the corresponding regions of the filtered widefield image in the second row. Subfigures a and b are the maximum intensity projections of a 10 µm thick z-stack obtained from 25 µm thick liver sections perfused with BioTracker NIR790 cytoplasmic membrane dye. The insets i and I are maximum intensity projections of the entire stack. The difference between OS-SIM and filtered widefield is clearly visible – much more structures and even small holes are visible in i that do not appear in I. The single z-slices ii and II show the stained nuclear membrane. Scale bar a,b: 10 µm; I,II,i,ii: 2 µm.*

### 2. Confocal imaging of a multicolor stained liver slice

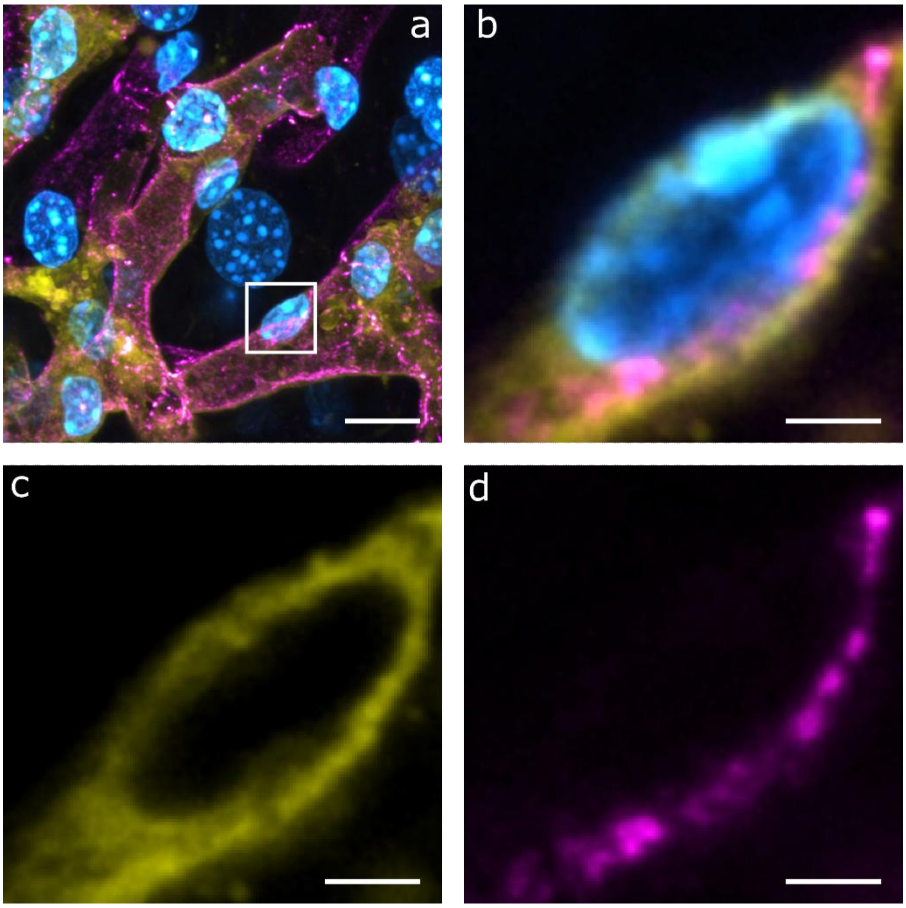


***Figure S3. Confocal image of a multicolor perfusion of the liver slice to verify the staining of the nuclear membrane.*** *The signal of PECAM-1 antibody is shown in magenta, the nuclear stain Hoechst 33258 is shown in cyan and the fluorescence signal of the BioTracker 555 dye is presented in yellow. A maximum intensity projection of a 25 µm thick liver slice with all three channels (a) visualizes the whole vessel network. The cropped region (b) is a single z-slice of the stack with an elogated nucleus of endothelial cells. The BioTracker 555 signal stains the nuclear membrane (c) and the PECAM-1 signal can only be detected at the inner cell surface. Scale bar a: 10 µm; b-d: 2 µm.*
